# α-Synuclein Overexpression and the Microbiome Shape the Gut and Brain Metabolome in Mice

**DOI:** 10.1101/2024.06.07.597975

**Authors:** Livia H. Morais, Joseph C. Boktor, Siamak MahmoudianDehkordi, Rima Kaddurah-Daouk, Sarkis K. Mazmanian

**Affiliations:** Division of Biology and Biological Engineering, California Institute of Technology, Pasadena, CA, USA; Aligning Science Across Parkinson’s (ASAP) Collaborative Research Network, Chevy Chase, MD 20815; Department of Psychiatry and Behavioral Sciences, Duke University, Durham, NC, USA; Duke Institute of Brain Sciences, Duke University, Durham, NC, USA; Department of Medicine, Duke University, Durham, NC, USA

**Author notes:** Corresponding Authors: Rima Kaddurah-Daouk, Sarkis K. Mazmanian email. Contributed equally (Livia H. Morais, Joseph C. Boktor, Siamak MahmoudianDehkordi).

## Abstract

Pathological forms of the protein α-synuclein contribute to a family of disorders termed synucleinopathies, which includes Parkinson’s disease (PD). Most cases of PD are believed to arise from gene-environment interactions. Microbiome composition is altered in PD, and gut bacteria are causal to symptoms and pathology in animal models. To explore how the microbiome may impact PD-associated genetic risks, we quantitatively profiled nearly 630 metabolites from 26 biochemical classes in the gut, plasma, and brain of α-synuclein-overexpressing (ASO) mice with or without microbiota. We observe tissue-specific changes driven by genotype, microbiome, and their interaction. Many differentially expressed metabolites in ASO mice are also dysregulated in human PD patients, including amine oxides, bile acids and indoles. Notably, levels of the microbial metabolite trimethylamine N-oxide (TMAO) strongly correlate from the gut to the plasma to the brain, identifying a product of gene-environment interactions that may influence PD-like outcomes in mice. TMAO is elevated in the blood and cerebral spinal fluid of PD patients. These findings uncover broad metabolomic changes that are influenced by the intersection of host genetics and the microbiome in a mouse model of PD.

## Introduction

Parkinson’s disease (PD) is the second most prevalent neurodegenerative condition, affecting 3% of the elderly population^1^ and presenting a significant social and economic burden that is growing as lifespans increase^2^. The hallmark symptom of PD is progressive movement dysfunction, which can include tremors, stiffness, and difficulty with balance and coordination. Currently available treatments can be effective but often induce undesirable side effects, are difficult to dose, and are not disease-modifying. The etiology of PD is multifactorial, with both genetic and environmental factors contributing to pathophysiology^3,4^. Mutations in, or overexpression of, the *SNCA* gene which encodes for the neuronal α-synuclein (αSyn) protein increases risk for developing PD^5^. In healthy neurons, αSyn regulates cellular homeostasis by modulating synaptic function and neurotransmitter release^6^. In pathological conditions, conformational changes in αSyn, phosphorylation at serine residue 129, and accumulation of aggregated forms are associated with synucleinopathies, a family of disorders with wide-ranging clinical presentations^7^, with PD being the most prevalent and best studied. Pathological species of αSyn can also be found outside the central nervous system (CNS). For example, phosphorylated αSyn is markedly elevated in the gastrointestinal (GI) tract of patients up to 20 years before PD diagnosis^8–10^, and many individuals with PD will experience clinical GI symptoms in the prodromal phase, with constipation being correlated to PD severity^11^. Braak et al. proposed that some forms of PD may originate in the gut and subsequently spread to the brain, potentially explaining why GI symptoms precede motor deficits^12^. In animal models, αSyn pathology can propagate via neurons from the gut to the brain^13,14^.

Alterations in the gut microbiome, known as dysbiosis, are observed in neurodegenerative disorders such as PD and their associated preclinical models^15–20^. The fecal microbiome in human PD patients, compared to matched controls, contains fewer anti-inflammatory bacteria (i.e., *Blautia, Coprococcus, Faecalibacterium*, and *Roseburia*), while pathogenic taxa (i.e., *Streptococcus*, *Enterococcus,* and *Actinomyces*) are increased in relative abundance^21–24^.

Functional analysis of the gut microbiome using shotgun metagenomics suggests that microbial metabolism and downstream metabolic pathways are altered in PD. For instance, a systematic analysis of the gut microbiome of individuals with PD revealed impaired bacterial amino acid synthesis, increased levels of homocysteine, and decreased levels of glutamate and glutamine^25^. Bacterial folate biosynthesis is also decreased in PD patients compared to healthy individuals^26^. In addition, the gut microbiome can alter the metabolism and systemic availability of the main drug used in the treatment of PD, levodopa (L-dopa)^20,27,28^. Metabolomic surveys of various biological samples (i.e., cerebrospinal fluid (CSF), plasma, serum, sebum, saliva, and feces) have uncovered changes in the presence and levels of various amino acids, amines, urate, lipids, and other chemicals in PD^29–34^. While gut bacteria are significant modulators of the metabolite repertoire in humans, contributing an estimated 50% of the small molecules in blood^35^, whether and how PD is impacted by metabolomic dysregulation remain unknown.

In mice, overexpression of human αSyn from a Thy1 promoter models certain forms of PD that may result from gene duplication or increased gene expression. αSyn-overexpressing (ASO) mice exhibit progressive motor deficits, GI symptoms, and αSyn pathology in the gut and brain^36,37^. Our laboratory has revealed that ASO mice raised in germ-free conditions or treated with broad spectrum antibiotics, i.e., mice without a microbiome, do not develop motor dysfunction and do not show αSyn aggregates in the brain^19^. We also showed that fecal microbiota transplant (FMT) from human PD patients into ASO mice worsens motor deficits compared to FMT from healthy donors^19^. Interestingly, infection of PD mouse models with enteric pathogens or induction of intestinal inflammation worsens PD-like phenotypes^38–41^. Conversely, dietary interventions that restore healthy microbiome profiles ameliorate motor deficits and αSyn pathology in the substantia nigra and striatum of ASO mice^42^. The microbiome is also altered in non-human primate models of PD^43,44^, and numerous microbiome surveys in humans have shown stereotypical changes in the fecal microbiome between PD patients and household and population controls^45–49^. Collectively, these studies support the hypothesis that the microbiome is an environmental modifier of genetic risk in PD. Based on the microbiome’s profound impacts on metabolism, we performed targeted metabolomic profiling of ASO mice under standard laboratory or germ-free housing conditions. We report tissue-specific metabolite changes driven by genetics, the microbiome, or both that are reminiscent of metabolomic signatures in human PD. We also identify characteristic changes in microbial molecules that link the gut and the brain. These findings advance our understanding of gene-environment interactions associated with synucleinopathies such as PD.

## Results

### Metabolomic profiles in mice are shaped by αSyn overexpression and the microbiome

To explore the effects of genotype and microbiome on the metabolome in a PD mouse model, “Line 61” Thy1-ASO and wild-type (WT) littermates were reared in specific pathogen-free (SPF; standard laboratory microbiota) or germ-free (GF) conditions to 4 months of age, when ASO mice display robust motor symptoms and constipation-like phenotypes^19^. Since the transgene is carried on the X chromosome, only male animals were used to avoid the effects of X-inactivation, standard practice in studies with this mouse model^37,50–53^. Samples were collected from colon tissue, colonic contents, cecal contents, duodenum, duodenal contents, plasma, brainstem, substantia nigra, striatum, and cortex, and analyzed by quantitative metabolomics. We employed the biocrates MxP Quant 500 platform, which measures 630 unique metabolites across 26 biochemical classes chosen to capture influences of diet and host-microbial interaction. We applied linear regression analysis incorporating covariates to correct for body weight, and t-distributed stochastic neighbor embedding (t-SNE) to reduce the dimensionality of metabolomic profiles for optimal visualization. Across all samples, we observed chemical feature clustering by tissue (**Fig. 1a**), as expected.

**Fig. 1:**
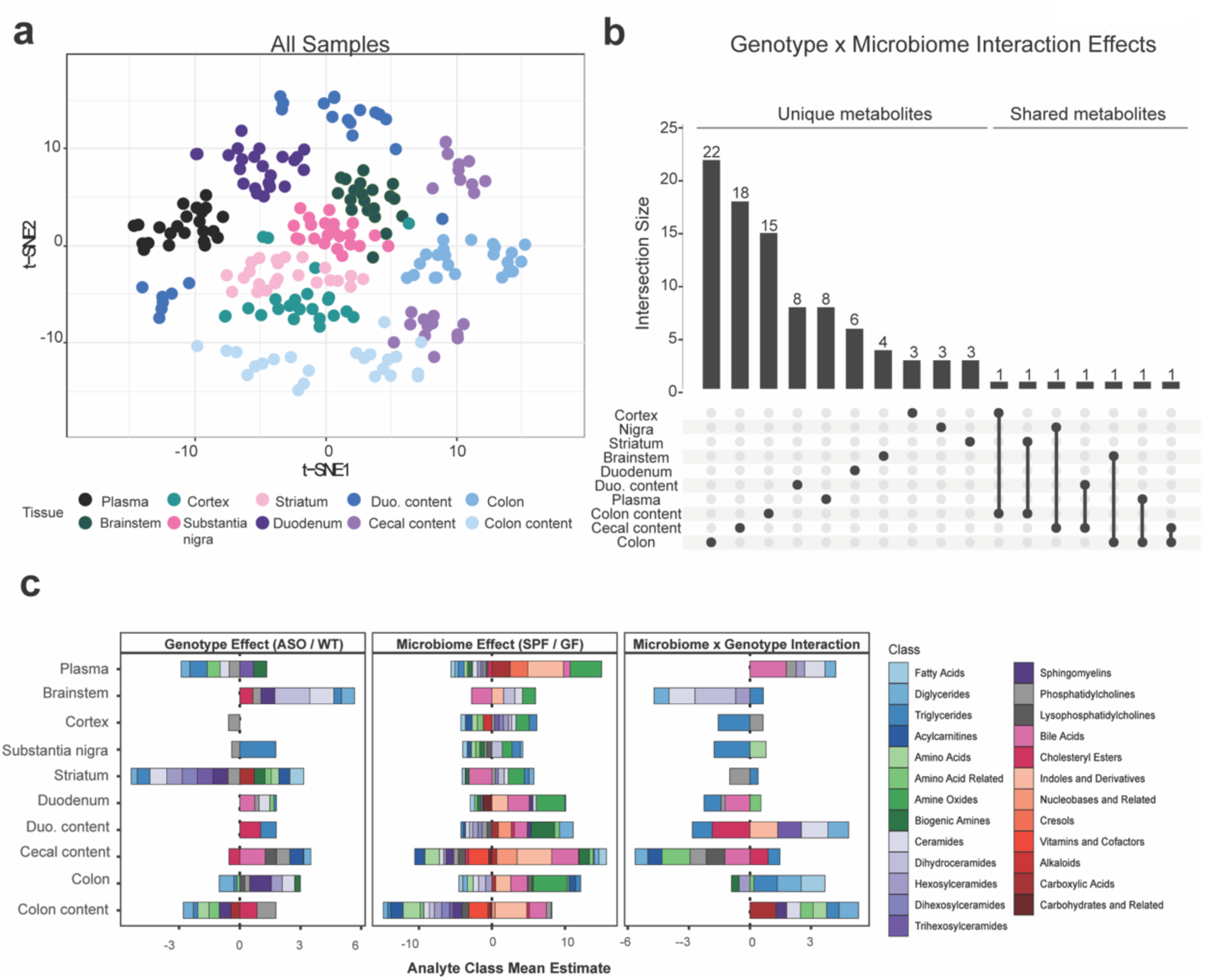
αSyn overexpression and microbiome presence alter global metabolomic profiles in mice. **a)** tSNE plot of all metabolomic samples in this study, colored by tissue. **b)** UpSet plot of unique and shared metabolite sets across all samples. Interaction size describes the number of metabolites with a significant genotype×microbiome interaction effect (p<0.05). The dots below the bar chart indicate the sample source of the metabolites. Singular dots with no vertical lines connecting to other tissues indicate a set of metabolites which are uniquely altered in a particular tissue. **c)** Stacked barplots displaying metabolite classes significantly associated (p<0.05) with either a genotype, microbiome, or genotype×microbiome interaction effect in a linear regression model.

In order to analyze conditional genotype (WT vs. ASO) and/or microbiome (SPF vs. GF) effects on the metabolome, we employed linear regression modeling that defined metabolites of interest that met either of the following criteria: 1) altered between ASO-SPF and WT-SPF animals, reflecting a genotype effect; and/or 2) altered between ASO-SPF and ASO-GF animals, reflecting a microbiome effect. Due to the large number of samples and the complexity of the experimental design, few metabolites passed a false discovery rate (FDR) threshold of 0.1; instead, we present all findings with *p* values ≤ 0.05 (**Table S1**). Across all tissues, we observed sets of metabolites significantly altered by genotype (**Fig. S1a**), microbiome (**Fig. S1b**), or the interaction between both (**Fig. 1b**). Individual metabolites displayed pronounced tissue specificity, with limited overlap between the gut and the brain (**Fig. 1b**). However, three metabolites impacted by the interaction between genotype and microbiome were shared between the gut and brain: triglyceride (TG) (16:0_40:7), diglyceride (DG) (14:1_18:1), and taurine. Additional molecules found in both tissues showed either genotype- or microbiome-specific changes (**Fig. S1**). We observed that genotype primarily influenced metabolites in the striatum, whereas the gut microbiome predominantly affected the metabolome in the colon, colonic contents, and cecal contents. Similarly, most of the analytes that represented interactions between and the genotype and microbiome were altered in the gut (**Fig. S1c**).

To assess functional changes in the metabolome, we grouped significantly altered metabolites by biochemical features (**Fig. 1c** and **Table S1**). We observed a broad range of molecular classes affected by the presence or absence of a microbiome, compared to more limited effects between ASO and WT animals. The microbiome influenced metabolite classes in all biological samples, with adjacent sites within gut or brain tissues displaying similar alterations (**Fig. 1c**). In SPF mice, we observed enrichment of metabolite classes synthesized and modulated by gut microbes, including amine oxides, bile acids, and indoles and their derivatives. In ASO mice, the most significant alterations occurred in lipid metabolites (**Fig. 1c**). At the molecular class level, there were minimal genotype-microbiome interactions.

### The gut microbiome modulates levels of bioactive metabolites in the gut of ASO mice

Having broadly identified classes of metabolites influenced by αSyn overexpression and the microbiome, we next focused on metabolomic profiles in the gut, where we observed the most pronounced metabolite changes in response to microbiome status. Microbial communities vary spatially along the gastrointestinal (GI) tract, and accordingly we analyzed the metabolome in different regions of the small and large intestines. At the level of functional metabolite classes, we discovered that the microbiome induced highly concordant shifts across all gut tissues, while genotypic effects were more specific to different segments of the gut (**Fig. 2a**), suggesting that αSyn overexpression and/or aggregation is not uniform along the GI tract in this model.

**Fig. 2:**
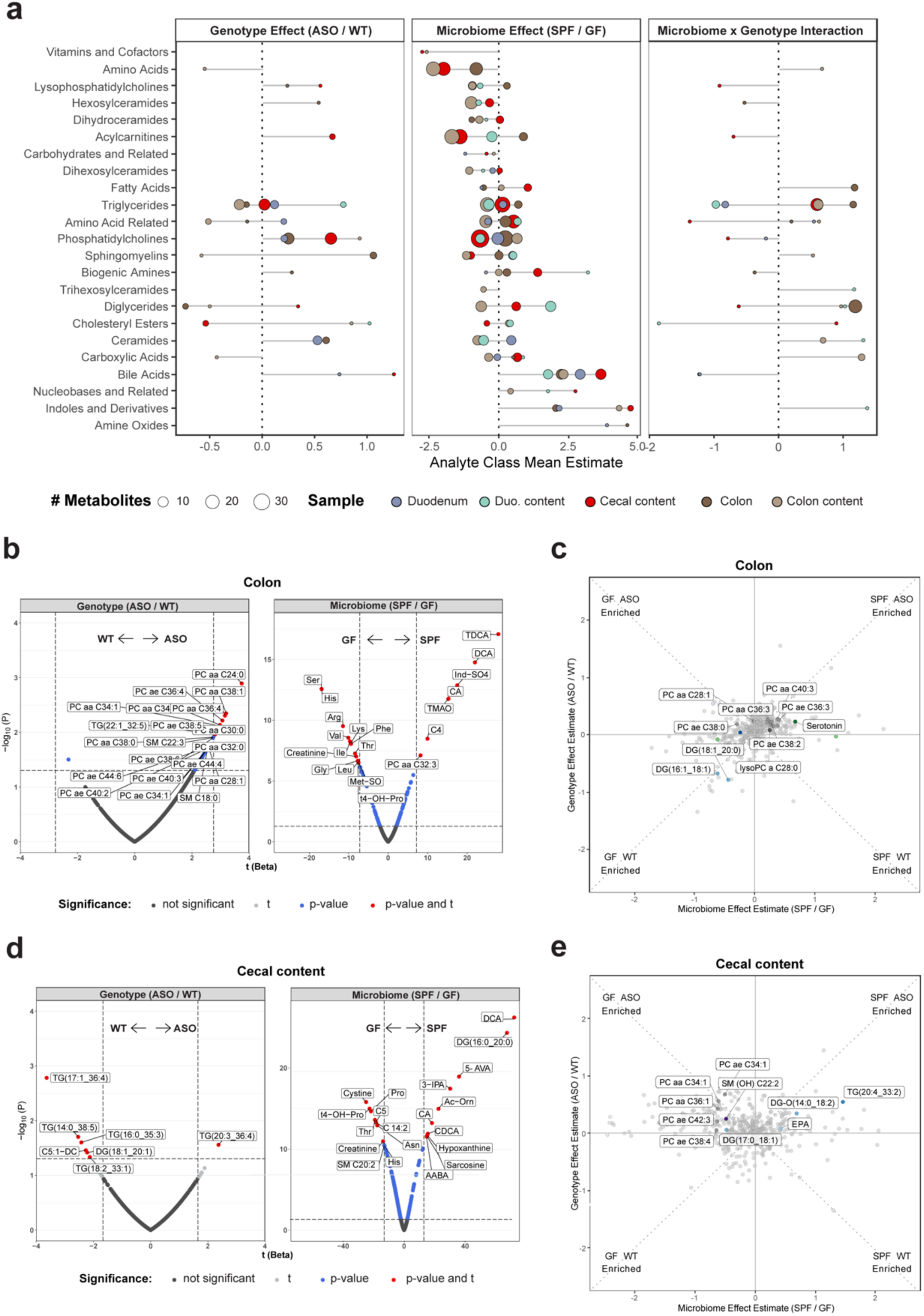
The gut microbiome shapes metabolism similarly across the GI tract. **a)** Lollipop plot of relative enrichment/depletion of significantly altered (p<0.05) metabolites in gut samples, organized by molecular class. Data points are colored by sample source and sized by number of metabolites. **b,d**) Volcano plots showing metabolites significantly enriched by genotype or microbiome status in the colon (**b**) and cecal contents (**d**). **c,e**) Scatterplots of metabolites in the colon (**c**) and cecal contents (**e**) affected by the microbiome and genotype in a linear model. Colored and labeled metabolites are the 25 metabolites showing the most significant (p<0.05) genotype×microbiome interaction effect.

Consistently altered metabolites in ASO vs. WT animals included TGs and DGs, as well as phosphatidylcholines (PC), the most abundant membrane phospholipid^54^. ASO mice displayed higher overall PC abundance in the colon **(Fig. 2b,c**), with elevated levels of PC aa C28:1 and PC ae C38:0 specifically in ASO-GF mice (**Fig. 2c**). A similar trend was evident in cecal contents, where PC aa C34:1 and PC aa C36:1 were more abundant in ASO-GF compared to ASO-SPF animals (**Fig. 2d,e**). In SPF vs. GF animals, we observed increased levels of nucleobases, and bile acids such as deoxycholic acid (DCA) and taurodeoxycholic acid (TDCA) (**Fig. 2, Fig. S2**). Bile acids are produced in the liver and conjugated by gut bacteria into secondary bile acids such as DCA and TDCA, which have emerged as important signaling molecules that regulate metabolic and signaling functions, including lipid absorption, cholesterol clearance, and nuclear receptor activation^55–57^. These microbial metabolites have diverse functions ranging from facilitating nutrient absorption to impacting immune responses in the gut and systemic compartments, and altered bile acid profiles in the brain have been associated with depression and Alzheimer’s disease (AD)^58–60^. We also discovered changes in levels of indoles and their derivatives, including indoxyl sulfate (Ind-SO_4_) (**Fig. 2, Fig. S2**). Ind-SO_4_ is a urinary metabolite that is elevated in anxiety and depression and has been shown to induce neuropsychiatric symptoms in animal models^61,62^. Collectively, these findings reveal that microbial metabolites associated with signaling to the immune and nervous systems are dysregulated in the gut of ASO mice.

### The metabolome in PD-relevant brain regions is differentially affected by the microbiome

We next explored genotype-microbiome interactions in shaping the brain metabolome. PD is primarily associated with the progressive loss of dopaminergic neurons in the substantia nigra that project to the striatum^63^. However, neurodegeneration and pathology also occur in other areas^63^, and accordingly we examined various brain regions, including the brainstem, cortex, substantia nigra, and striatum. We observed clustering of metabolomes within each brain tissue and less separation between brain tissues compared to samples from the gut (**Fig. 1a**), indicating a brain-specific metabolomic signature. Comparing ASO to WT animals, we report unique differences in the metabolome within the striatum (**Fig. 3a**), with a notable increase in neuroactive amino acids—anserine, creatinine, and aconitic acid (AconAcid) (**Fig. 3b,c**). Interestingly, anserine levels were further enriched in ASO-GF animals, and proline (Pro), another oxidative stress modulator^64^, was higher in ASO-SPF mice than in other animal groups (**Fig. 3c**). ASO mice also harbored higher levels of phenylalanine (Phe) and tryptophan (Trp), precursors of dopamine and serotonin, respectively, in the striatum (**Fig. 3b**). In the cortex, ASO mice contained elevated levels of 3-methylhistidine (3-Met-His) (**Fig. S3c**). ASO mice showed decreased abundance of several lipids, including ceramides, TGs, and PCs throughout the brain, but particularly in the striatum and cortex (**Fig. 3b,c** and **Fig. S3).** Interestingly, a unique converse effect was observed in the brainstem, with more TGs in ASO mice (**Fig. S3a**). The brainstem, which is innervated by the autonomic nervous system, serves as a central hub for lipid sensing^65^, suggesting potential systemic dysregulation of lipid metabolism in response to αSyn overexpression.

**Fig. 3:**
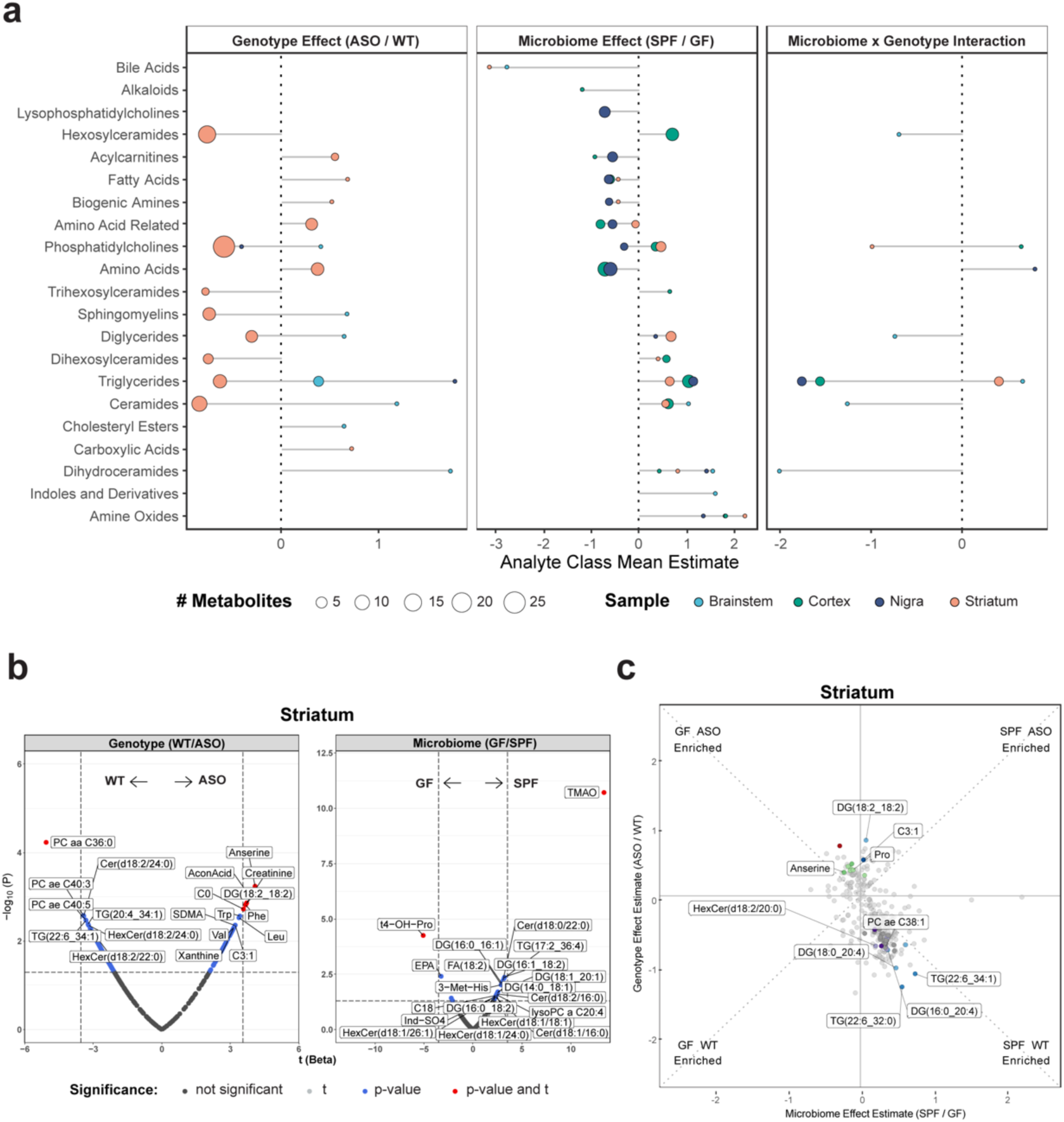
The genotype and microbiome alter metabolite levels differentially across the brain. **a**) Lollipop plot of relative enrichment/depletion of metabolites whose levels are significantly altered (p<0.05) by genotype, microbiome, or their interaction in the brain. Data points are colored by tissue and sized by number of metabolites. **b**) Volcano plots showing metabolites significantly enriched by genotype or microbiome status in the striatum. **c**) Scatterplot of metabolites in the striatum affected by the microbiome and genotype in a linear model. Colored and labeled metabolites are the 25 metabolites showing the most significant (p<0.05) genotype×microbiome interaction effect.

Changes in the metabolome were evident across all brain regions (**Figs. 3**, **S3**). Regardless of genotype, the gut microbiome significantly shaped lipid metabolism, influencing key metabolites such as TGs, DGs, ceramides, and lysophosphatidylcholine (LysoPC), with notable enrichment of lipids in the brains of SPF animals (**Figs. 3, S3)**. The most dramatic influence of the microbiome throughout the brain was a significant increase in levels of trimethylamine N-oxide (TMAO) (**Figs. 3b, S3**).

### A single microbially-synthesized metabolite connects the GI tract, plasma, and the brain

To explore links between altered central and peripheral metabolism, we examined the plasma metabolome, which serves as a surrogate for chemical communication between the gut and the brain. In ASO animals, we observed moderate depletion of lipid metabolites (**Fig. 4a**), suggestive of dysregulation in lipid homeostasis, with DGs, TGs, and PCs being the most affected by the microbiome (**Fig. 4b**). Valine, a branched-chain amino acid (BCAA) involved in energy generation^66^, was enriched in GF animals compared to SPF counterparts, highlighting the influence of gut bacteria on amino acid metabolism (**Fig. 4c**).

**Fig. 4:**
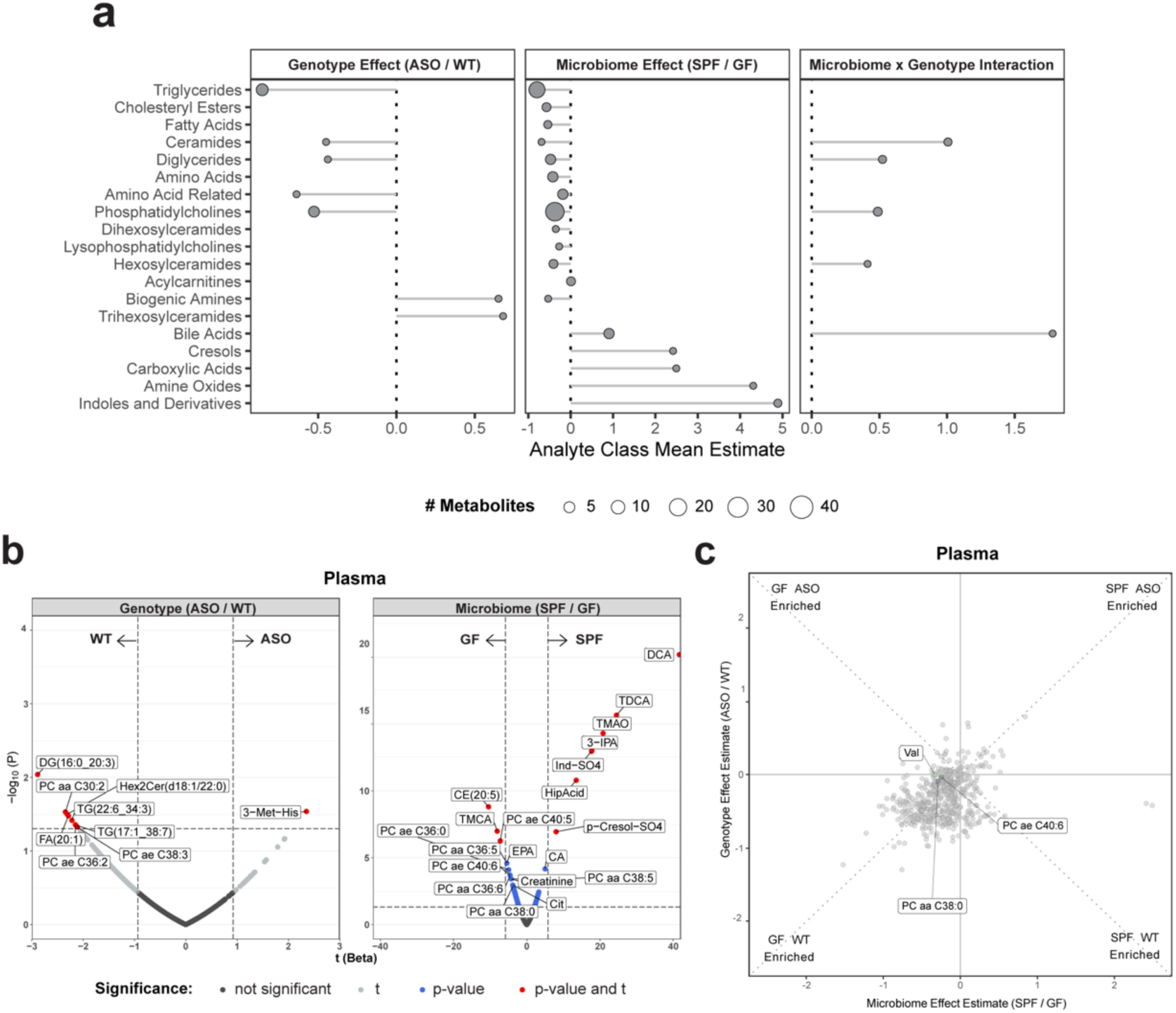
αSyn overexpression and microbiome influence circulating metabolites. **a**) Lollipop plot of relative enrichment/depletion of metabolites whose levels are significantly altered (p<0.05) by genotype, microbiome, or their interaction in plasma. Data points are colored by metabolite class and sized by number of metabolites. **b**) Volcano plots showing metabolites significantly enriched by genotype or microbiome status in the plasma. **c**) Scatterplot of metabolites in the plasma affected by the microbiome and genotype in a linear model. Colored and labeled metabolites are the 25 metabolites showing the most significant (p<0.05) genotype×microbiome interaction effect.

When we correlated levels of metabolites detected in plasma with their levels in other tissues, molecular features with the strongest plasma level correlation were those linked to gut microbes (**Fig. 5**). In the brain, these metabolites included Ind-SO_4_, amino acid-related compounds, and TMAO. The levels of many additional metabolites were strongly correlated between the plasma and gut tissues and contents, including bile acids, indoles and their derivatives, and amino acid-related molecules. TMAO was notable in being strongly correlated between the GI tract and plasma, as well as between plasma and brain tissues (**Fig. 5**). TMAO has been widely studied in a number of microbiome-related conditions and elevated levels are associated with cardiovascular disease, chronic kidney disease, metabolic disorders, GI cancers and stroke^67–72^. Importantly, TMAO levels are increased in the plasma and cerebrospinal fluid (CSF) of PD patients relative to controls^73,74^ and gene families involved in TMA production are elevated in the PD gut microbiome^24^, though some conflicting results have been reported^75,76^. Our study design, analyzing multiple sites from the gut to the brain, uncovers TMAO as a potentially pathogenic molecule that may link the gut microbiome to the brain in ASO mice.

**Fig. 5:**
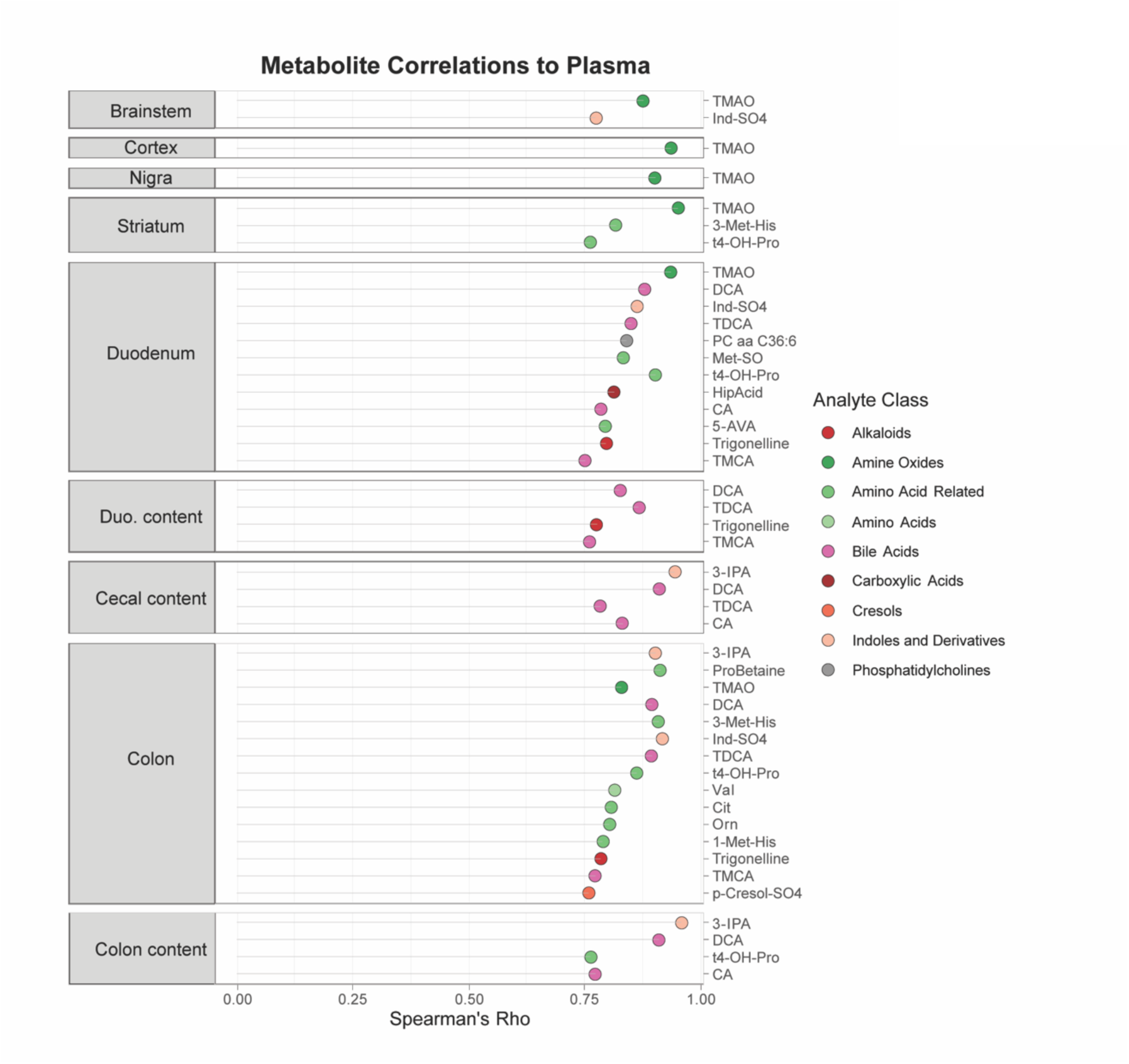
A microbially-produced metabolite links the gut-brain axis in ASO mice. Lollipop plot showing metabolites whose levels in each indicated sample correlate strongly (Spearman’s Rho > 0.75) with the level of the same metabolite in plasma.

## Discussion

αSyn overexpression (a genetic factor) and microbiome alterations (an environmental factor) have been implicated in PD. We reveal herein that gene-microbiome interactions shape the metabolome in a mouse model of synucleinopathy. Notably, many of the metabolites that were altered in ASO mice have been implicated in human PD, affecting mitochondrial function, oxidative stress response, inflammation, protein aggregation, and neurotransmission. As the microbiome is altered in PD patients, and gut bacteria impact motor and non-motor symptoms in several PD models, dysregulation of microbial metabolites may represent a key aspect of PD pathophysiology that warrants further investigation.

Lipid homeostasis has been proposed as a key feature of PD^77^. We uncovered that both genotype and the microbiome influence lipid abundance, particularly of PCs in the brain and gut.

Alterations in lipid metabolism have been widely implicated in PD and other neurodegenerative disorders, including Alzheimer’s disease^78,79^. Interestingly, PCs influence the aggregation of αSyn^80^. Brains of individuals with PD display decreased PC in the visual cortex^81^, and a similar reduction was seen in a rat model of early-stage PD^82^. In the gut, PC plays a significant role in the colonic mucosa, forming a hydrophobic barrier to protect from inflammatory insults^83^. Lower levels of PCs in the luminal mucus may contribute to gut mucosal inflammation and other GI issues such as constipation and delayed gastric emptying in PD^84^. In addition to reduced levels in the gut and brain, we observed moderate depletion of lipid metabolites, particularly PCs and TGs, in the plasma of ASO mice, suggesting widespread dysregulation of lipid homeostasis. In humans, an elevated PC/lysophosphatidylcholine (lyso-PC) ratio has been reported in the plasma of individuals with PD^85^. Similarly, lower levels of serum TGs, non-esterified fatty acids, and cholesterol are observed in the A53T human αSyn mouse model and correlate with weight loss^86^. In humans, lower serum and plasma levels of TGs and cholesterol are linked to PD^87–89^, and alterations in lipid metabolism have also been described in AD and other neurodegenerative conditions^90–92^.

In gut-derived samples, we found interesting profiles for microbially-derived metabolites implicated in inflammation. As expected, GF mice have lower levels of secondary bile acids and indole metabolites, including Ind-SO_4_. Altered bile acid production has been observed in the GI tracts of individuals with PD^47,93^ as well as in αSyn -based mouse models^94^. Elevated levels of secondary bile acids such as DCA and lithocholic acid (LCA) are associated with the prodromal stage of PD and are linked to an increase in Clostridiales cluster XI in the gut^94^. Functional microbiome analysis and metabolic modeling has suggested that *Akkermansia muciniphila*, *Arcanobacterium ihumii*, *Alistipes shahii*, and *Candidatus* Gastranaerophilales increase production of indole and its derivatives such as indole*-*3*-*propionic acid (IPA), resulting in elevated levels in the serum of PD patients^95^. Another indole metabolite, Ind-SO_4,_ has been linked to increased oxidative stress^96^, a notable feature of PD pathology. The local effects of Ind-SO_4_ in the gut remain uncharacterized, but recent research suggests effects in the CNS, driving glial activation, neurodegeneration, anxiety, and cognitive deficits^97–99^. Remarkably, the concentration of Ind-SO_4_ in the urine of PD patients is doubled compared to individuals without _PD100._

In the brain, changes in amino acid profiles suggest effects on oxidative stress and neurotransmission, as well as energy metabolism. We found that ASO mice exhibit higher levels of neuroactive amino acids with crucial roles in regulating oxidative stress and redox homeostasis^101^. A recent study in the αSyn A53T mouse model of PD also observed accumulation of carnosine, homocarnosine, and anserine in the brain, which was theorized could be a defense against reactive oxygen species (ROS)^102^. Interestingly, we reveal that anserine levels were further enriched in ASO-GF animals, indicating a combinatorial effect of genotype and the microbiome. αSyn overexpression also increased levels of proline, an oxidative stress modulator^64^ that is increased in the serum of PD patients^103^, and AconAcid, a fatty acid in the tricarboxylic acid (TCA) cycle which also influences oxidative phosphorylation. In individuals with PD, the levels of AconAcid and other TCA cycle metabolites are reduced^104,105^; therefore it would be interesting to measure these molecules longitudinally in humans. Levels of Phe and Trp, precursors for dopamine and serotonin, respectively, were increased in the striatum of ASO mice. These neurotransmitter pathways are impacted in PD^106,107^, and our observations align with metabolomic profiling of CSF from PD patients showing that Phe and Trp accumulation correlates with disease progression^108^. Interestingly, among the brain regions profiled in our study, the striatum was the most metabolically imprinted by both genotype and the microbiome.

In the plasma, we uncovered elevated levels of valine, a BCAA, in ASO mice lacking a microbiome. A recent study identified changes in BCAA concentration in the serum of individuals with PD that correlated with disease stage and gut microbiome composition^87^. Furthermore, serum BCAA concentrations are lower in individuals with advanced-stage PD^87^. In dopaminergic neurons derived from stem cells from human PD patients, the relative abundance of valine is also diminished and is linked to disrupted mitochondria-lysosome contact dynamics^109^.

We modeled the flux of metabolites from the gut to the brain via plasma, and discovered a strong signal for the microbial metabolite TMAO. Trimethylamine (TMA) is produced by gut bacteria through conversion of dietary precursors such as choline, betaine, and L-carnitine, followed by conversion into TMAO in the liver^68^. TMAO has been associated with infiltration of inflammatory cells in the colon and rectal epithelium, as well as cellular damage^110^. TMAO is strongly implicated in arteriosclerotic cardiovascular disorders and systemic inflammation^111^. Importantly, levels of TMAO and related metabolites are increased in CSF and plasma from PD patients^112^. The role of TMAO in the CNS remains largely unclear, with both beneficial and detrimental effects reported^113,114^. TMAO is associated with brain inflammation, astrocyte activation, and cognitive deficits in mice^115^, as well as neuronal senescence and mitochondrial dysfunction^113^. In individuals with PD, increased levels of TMAO and other bacterial-derived metabolites have been implicated in cognitive decline^116^, and higher levels of TMAO in the serum and CSF correlate with disease severity and the progression of motor symptoms^117^. TMAO has been shown to induce fibrillar aggregation of αSyn in a concentration-dependent manner *in vitro*^118–120^. Our findings add to growing evidence supporting a potential role for TMAO in PD.

The metabolome serves as a comprehensive indicator of environmental influences, including contributions of diet, toxins, drugs, and gut microbiome. Herein, we reveal broad changes in the metabolomic profile in gut, plasma, and brain of a mouse model of αSyn overexpression. Our study is limited by sample size, use of a single mouse model, and exclusion of sex as a variable due to the necessity of performing experiments exclusively in male ASO mice. However, our use of GF mice allows unequivocal assignment of metabolites that are regulated or produced by the gut microbiota, which is not possible in human studies. Discovery of changes in specific microbial molecules across multiple tissues in ASO mice, many of which correlate to findings in human studies, highlights the need for future investigations into the mechanistic role of gut bacteria in the pathophysiology of PD and other synucleinopathies.

## Methods

### Mice

Male mice overexpressing human αSyn under the Thy1 promoter (“Line 61” Thy1-α-Synuclein, ASO) and WT mice were generated by crossing wild-type BDF1 males (Charles River, RRID:IMSR_CRL:099) with ASO heterozygous females^121^. This study used the following numbers of mice: n=5 WT-SPF; n=6 ASO-SPF; n=6 WT-GF; n=6-ASO-GF. GF mice were generated by caesarean section, with the offspring fostered by GF Swiss-Webster dams and maintained microbiologically sterile inside flexible film isolators^14,19,122^. SPF mice were housed in autoclaved micro-isolator cages. GF status was confirmed on a bi-weekly basis through 16S rRNA PCR of fecal-derived DNA and plating of fecal pellets on Brucella blood agar under anaerobic conditions and tryptic soy blood agar under aerobic conditions. Mice received food and water ad libitum, were maintained on the same 12-hour light-dark cycle and housed in the same facility. All animal husbandry and experiments were approved by the California Institute of Technology’s Institutional Animal Care and Use Committee (IACUC).

### Tissue dissection

At 4 months of age, mice were sacrificed by decapitation. The brain was rapidly removed and placed in an ice-chilled stainless steel coronal matrix. Brain tissue was sectioned in slices of approximately 1 mm. Substantia nigra, striatum, motor cortex (referred to as cortex) and caudal brainstem (referred to as brainstem) were dissected within three minutes using reference brain atlas coordinates^123^. Gut tissue and contents were dissected on an ice-chilled stainless steel dissection tray. All samples were weighed, snap-frozen in dry ice and stored at -80^°^C until processing. For protocol see dx.doi.org/10.17504/protocols.io.14egn3pkzl5d/v1.

### Plasma collection

Trunk blood was collected in EDTA-coated tubes and kept at room temperature before plasma separation. Plasma was separated by centrifugation at 2,500 ×g for 10 minutes. Plasma was transferred to a pre-cooled collection vial and stored at -80^°^C until processing. For protocol see dx.doi.org/10.17504/protocols.io.14egn3pkzl5d/v1.

### Quantitative targeted metabolomics

#### Sample Preparation

Samples were prepared using the MxP Quant 500 kit (biocrates life sciences AG, Innsbruck, Austria) in strict accordance with the manufacturer’s protocol. Plasma samples were centrifuged, and the supernatant was used for analysis. Brain, colon, and duodenum tissue samples were first suspended in 3 μL ethanol/phosphate buffer per mg tissue wet weight. The samples were then sonicated, vortexed and homogenized using a Precellys-24 instrument (Bertin Technologies, Montigny le Bretonneux, France), and the supernatant was used for analysis. For measurement of some metabolites, it was necessary to dilute the duodenum tissue samples 1:5 in buffer before the samples were centrifuged and the supernatant used for analysis. To extract metabolites from duodenal, cecal and colonic contents, samples were resuspended in extraction buffer (85% ethanol in phosphate buffer) and vortexed thoroughly until dissolved. After homogenization, the samples were ultrasonicated in a chilled bath for 5 min. Samples were then centrifuged and the supernatant was used for analysis. An additional 1:1,000 dilution was prepared for the analysis of highly concentrated bile acids. For protocol see: dx.doi.org/10.17504/protocols.io.261ge5pwyg47/v1.

#### Measurement

A mass spectrometry (MS)-based targeted metabolomics approach was used to determine the concentration of endogenous metabolites in a total of 226 samples: 23 plasma samples, 69 gut content samples (23 cecal contents, 23 colon contents, and 23 duodenum contents), and 134 tissue samples (88 brain, 23 colon, and 23 duodenum). Metabolites were quantified using the commercially available MxP^®^ Quant 500 kit (biocrates). The kit provides measurements of up to 634 metabolites across 26 biochemical classes. Lipids (e.g, acylcarnitines, glycerophospholipids, sphingolipids, triglycerides) and hexoses were measured by flow injection analysis-tandem MS (FIA-MS/MS) using a 5500 QTRAP® instrument (AB Sciex, Darmstadt, Germany) with an electrospray ionization (ESI) source for the plasma and tissue samples, and a Xevo TQ-S (Waters, Vienna, Austria) instrument with an ESI source for the gut content samples. Small molecules were measured by liquid chromatography-tandem MS (LC-MS/MS), also using a 5500 QTRAP® instrument for all samples. Gut tissue and content samples were also measured by LC-MS/MS on a Xevo TQ-S instrument. To quantitatively analyze metabolite profiles in the samples, a 96-well-based sample preparation device was used which consists of inserts that have been impregnated with internal standards. A predefined sample amount was added to the inserts. Next, a phenyl isothiocyanate (PITC) solution was added to derivatize some of the analytes (e.g., amino acids), and after the derivatization was complete, the target analytes were extracted with an organic solvent, followed by a dilution step. The obtained extracts were then analyzed by FIA-MS/MS and LC-MS/MS methods using multiple reaction monitoring (MRM) to detect the analytes. Data were quantified using appropriate mass spectrometry software, either Sciex Analyst® (https://sciex.com/, RRID:SCR_023651) or Waters MassLynx™, https://www.waters.com/waters/en_US/MassLynx-Mass-Spectrometry-Software-/nav.htm?cid=513164, RRID:SCR_014271), and imported into Biocrates MetIDQ™ software for further analysis. For protocol see: dx.doi.org/10.17504/protocols.io.261ge5pwyg47/v1.

#### Quality control

The raw Q500 metabolomic profiles included measurements of 634 metabolites in 226 samples. Quality control steps were performed as in prior publications^124,125^. Separately for each material type, metabolites with >30% of measurements above the lower limit of detection (LOD) in SPF animals were included (n= 539, 459, 503, and 297 remaining metabolites in plasma, GI tissue, gut content, and brain tissue, respectively). Imputation of <LOD values was performed using each metabolite’s LOD/2 value to increase statistical power. Since all samples for each material type were measured on one plate, no batch effect removal procedure was conducted. Metabolite concentrations were log_2_ transformed to achieve normal distribution. For protocol see: dx.doi.org/10.17504/protocols.io.261ge5pwyg47/v1.

#### Statistical analysis

All statistical analyses were conducted in R 4.2.2 (https://www.r-project.org/, RRID:SCR_001905). t-distributed stochastic neighbor embedding (t-SNE) dimensionality reduction scatterplots were generated of log_2_-transformed metabolite concentrations using the Rtsne (v0.1-3.1, https://github.com/jkrijthe/Rtsne, RRID:SCR_016342) package with default parameters. To evaluate the overall effect of genotype, microbiome and their interaction on metabolome in each sample type, Permutational Multivariate Analysis of Variance (PERMANOVA) was conducted using the “adonis2” function in the vegan package (http://cran.r-project.org/web/packages/vegan/index.html, RRID:SCR_011950) with 10,000 permutations and Euclidean distance. Multivariate homogeneity of group dispersions was tested using the “betadisper” and “permutest” functions in the vegan package. To investigate the effects of genotype and microbiome status on the metabolome, separate linear regression models were constructed for each sample. These models defined log_2_ metabolite concentration as the dependent variable, and body weight, microbiome, genotype and their interaction as independent variables. This allowed us to estimate the effects of genotype and microbiome status while controlling for variability in body weight. The linear model was specified as: (log_2_(metabolite [µM]) ∼ Genotype + Microbiome + Genotype*Microbiome + Body-weight). For a non-parametric measure of metabolite-metabolite associations, we performed Spearman’s rank-based correlation. For protocol see: dx.doi.org/10.17504/protocols.io.261ge5pwyg47/v1.

## Data Availability

The datasets generated and analyzed in this study are available from: https://zenodo.org/records/10841477.

## Code Availability

The underlying code for this study is available in GitHub and can be accessed via https://zenodo.org/records/10841477.

## Acknowledgments

We thank Dr. Gil Sharon for help with tissue dissections, Taren Thron for animal breeding, Yvette Garcia-Flores for logistical assistance, and Dr. Catherine Oikonomou for critical review of the manuscript. Metabolomics data collection, preprocessing, analysis, and interpretation were provided by the Alzheimer’s Gut Microbiome Project (AGMP) and the Alzheimer’s Disease Metabolomics Consortium (ADMC), funded wholly or in part by the following grants and supplements: R01AG046171, RF1AG051550, RF1AG057452, R01AG059093, U19AG063744, 3U19AG063744-04S1, RF1AG058942, U01AG061359, R01MH108348, R01AG081322 and FNIH: #DAOU16AMPA awarded to R.K.M. at Duke University in partnership with a large number of academic institutions. As such, the investigators within the AGMP and the ADMC, not listed specifically in this publication’s author’s list, provided data along with its pre-processing and prepared it for analysis, but did not participate in analysis or writing of this manuscript. A listing of AGMP Investigators can be found at https://alzheimergut.org/meet-the-team/. A complete listing of ADMC investigators can be found at: https://sites.duke.edu/adnimetab/team/. L.H.M. was partially supported by an American Parkinson’s Disease Association postdoctoral fellowship during the study. This research was funded in part by Aligning Science Across Parkinson’s (ASAP-020495 and ASAP-000375 ) through the Michael J. Fox Foundation for Parkinson’s Research (MJFF) to S.K.M. For the purpose of open access, the authors have applied a CC BY public copyright license to all Author Accepted Manuscripts arising from this submission.

## Author Contributions

L.H.M. and S.K.M. conceived and designed the study. L.H.M. performed animal work and collected samples. S.M.D. performed metabolomics. S.M.D. and J.C.B. analyzed data and generated visualizations. S.K.M. and R.K.D. supervised the study. L.H.M. and S.K.M. wrote the manuscript, with input from all authors. All authors read and approved the final manuscript.

## Supplementary Figures

**Fig. S1:**
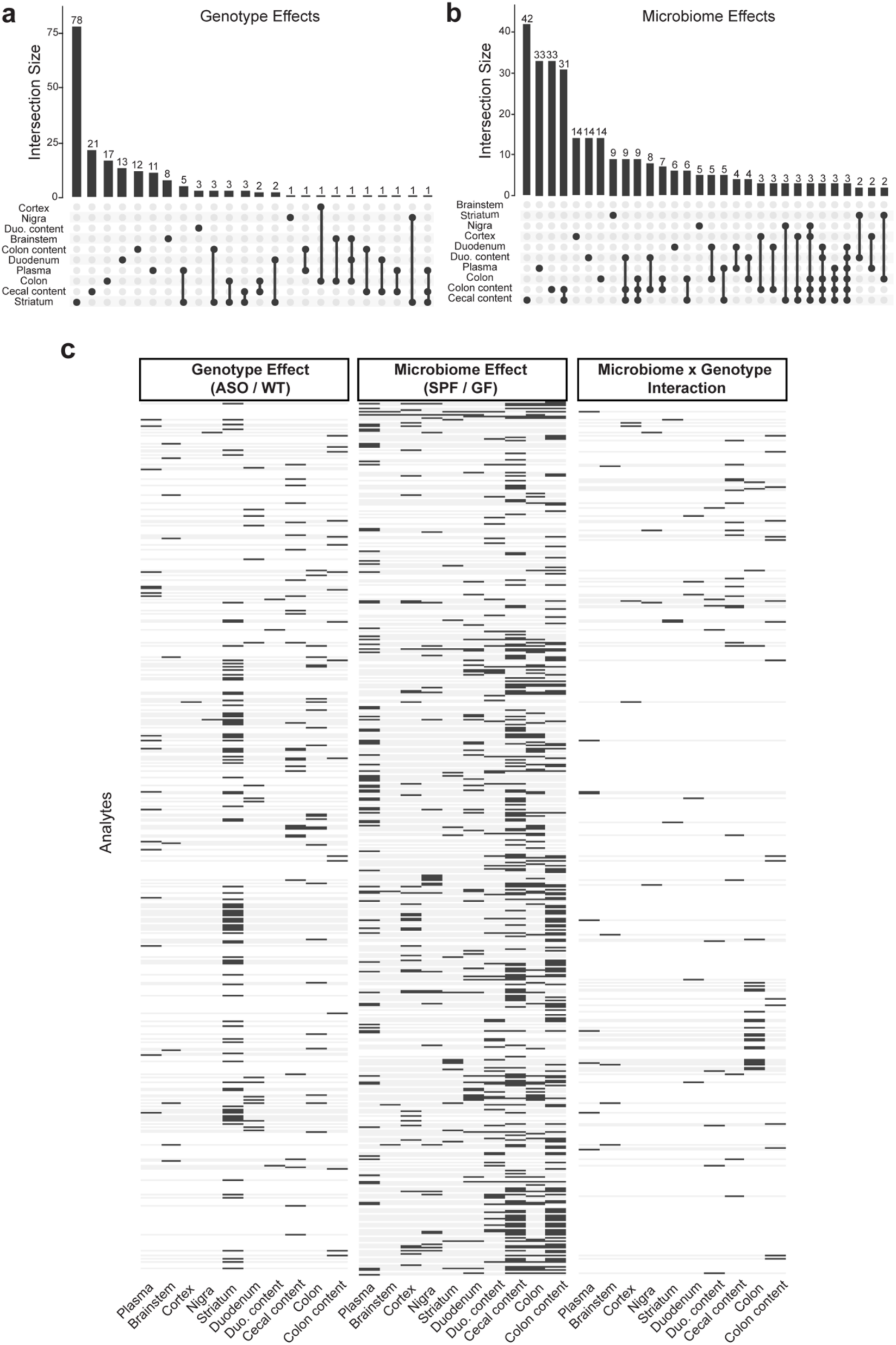
**a-b)** UpSet plots of unique and shared metabolite sets across all samples. Interaction size describes the number of metabolites with a significant genotype (**a**) or microbiome (**b**) interaction effect (p<0.05). The dots below the bar chart indicate the tissue source of the metabolites. Singular points with no vertical lines connecting to other tissues indicate a set of metabolites which are uniquely altered in a particular tissue. **c**) Binary heatmap of metabolites significantly altered in at least one tissue for at least one variable (genotype, microbiome, microbiome×genotype) from the linear model.

**Fig. S2:**
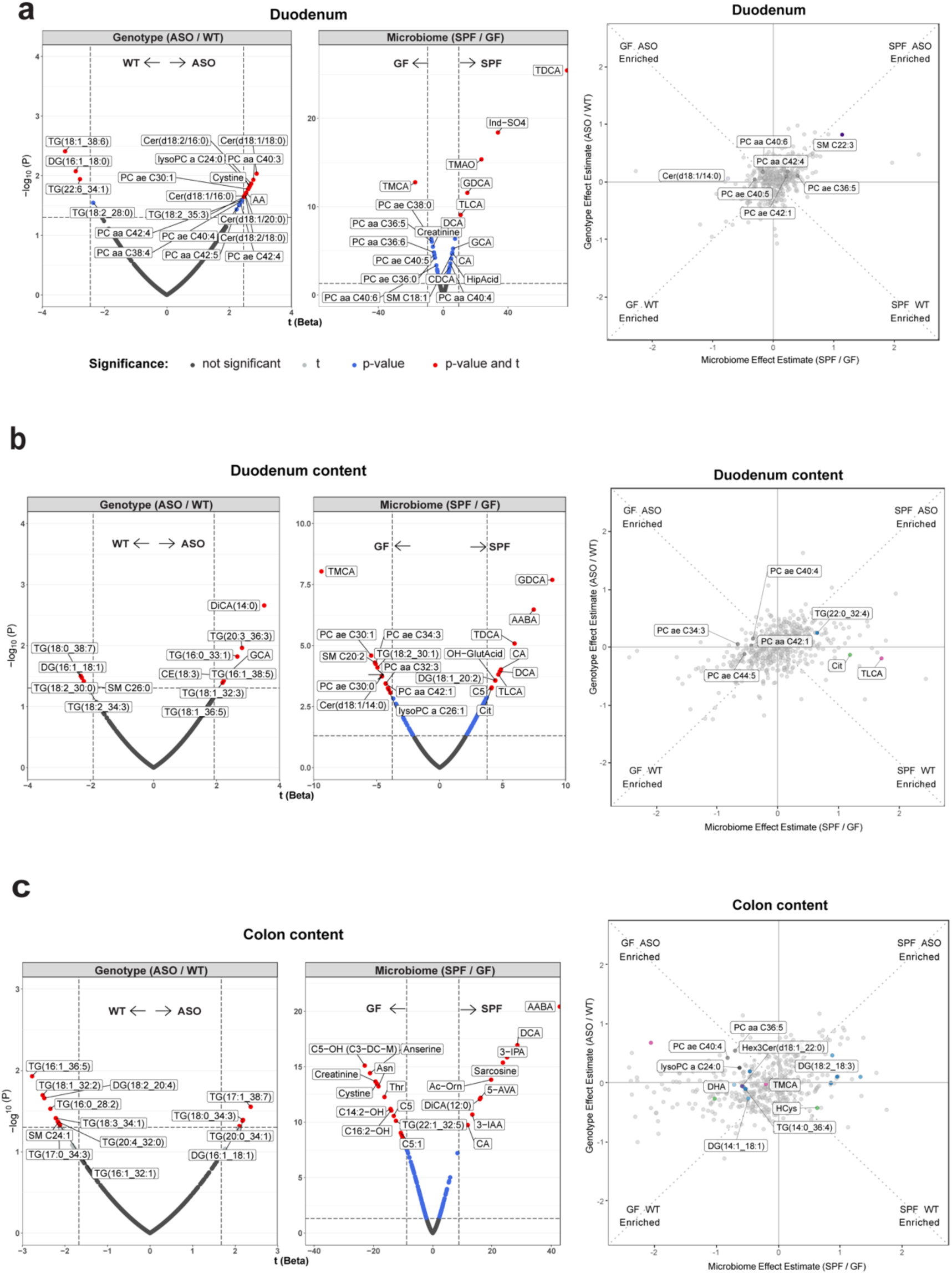
Volcano plots and scatterplots of the most significantly altered metabolites showing genotype, microbiome, or genotype×microbiome interaction effects in the duodenum (**a**), duodenal contents (**b**), or colonic contents (**c**). Conventions are as in Fig. 2.

**Fig. S3:**
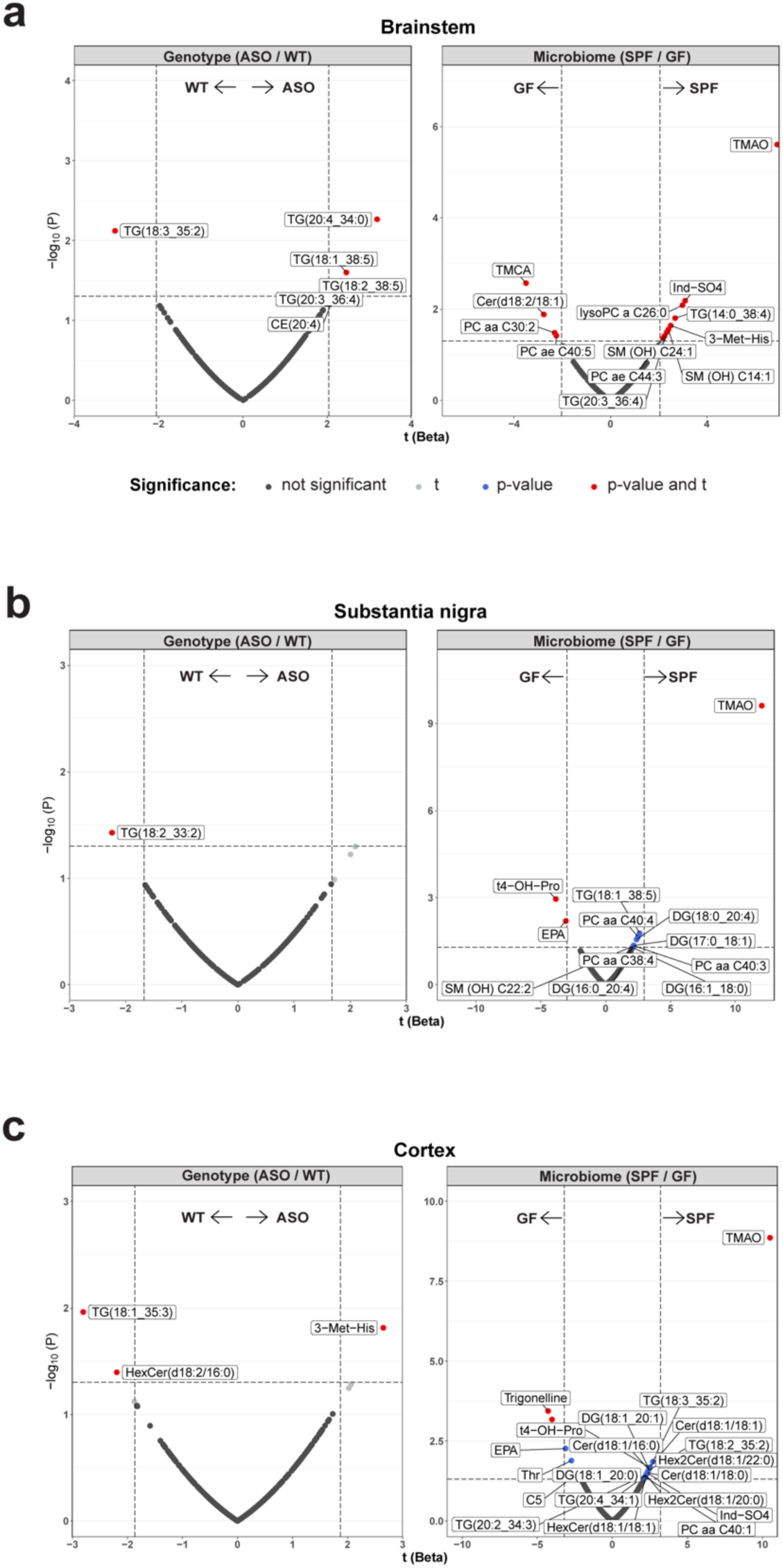
Volcano plots of metabolites showing genotype or microbiome effects in the brainstem (**a**), substantia nigra (**b**) and cortex (**c**).

## References

1. Aarsland, D. et al. Parkinson disease-associated cognitive impairment. Nat Rev Dis Primers 7, 1–21 (2021).

2. Yang, W. et al. Current and projected future economic burden of Parkinson’s disease in the U.S. npj Parkinsons Dis. 6, 1–9 (2020).

3. Klein, C. & Westenberger, A. Genetics of Parkinson’s Disease. Cold Spring Harb Perspect Med 2, a008888 (2012).

4. Ascherio, A. & Schwarzschild, M. A. The epidemiology of Parkinson’s disease: risk factors and prevention. The Lancet Neurology 15, 1257–1272 (2016).

5. Polymeropoulos, M. H. et al. Mutation in the α-Synuclein Gene Identified in Families with Parkinson’s Disease. Science 276, 2045–2047 (1997).

6. Somayaji, M. et al. A dual role for α-synuclein in facilitation and depression of dopamine release from substantia nigra neurons in vivo. Proceedings of the National Academy of Sciences 117, 32701–32710 (2020).

7. Houser, M. C. & Tansey, M. G. The gut-brain axis: is intestinal inflammation a silent driver of Parkinson’s disease pathogenesis? npj Parkinson’s Disease 3, 3 (2017).

8. Jones, J. D. et al. A bidirectional relationship between anxiety, depression and gastrointestinal symptoms in Parkinson’s disease. Clinical Parkinsonism & Related Disorders 5, 100104 (2021).

9. Zinnen, A. D. et al. Alpha-synuclein and tau are abundantly expressed in the ENS of the human appendix and monkey cecum. PLOS ONE 17, e0269190 (2022).

10. Ruffmann, C. et al. Detection of alpha-synuclein conformational variants from gastro-intestinal biopsy tissue as a potential biomarker for Parkinson’s disease. Neuropathology and Applied Neurobiology 44, 722–736 (2018).

11. Sung, H.-Y., Park, J.-W. & Kim, J.-S. The Frequency and Severity of Gastrointestinal Symptoms in Patients with Early Parkinson’s Disease. JMD 7, 7–12 (2014).

12. Braak, H. et al. Staging of brain pathology related to sporadic Parkinson’s disease. Neurobiology of Aging 24, 197–211 (2003).

13. Kim, S. et al. Transneuronal Propagation of Pathologic α-Synuclein from the Gut to the Brain Models Parkinson’s Disease. Neuron 103, 627–641.e7 (2019).

14. Challis, C. et al. Gut-seeded α-synuclein fibrils promote gut dysfunction and brain pathology specifically in aged mice. Nat Neurosci 23, 327–336 (2020).

15. Needham, B. D., Kaddurah-Daouk, R. & Mazmanian, S. K. Gut microbial molecules in behavioural and neurodegenerative conditions. Nat Rev Neurosci 21, 717–731 (2020).

16. Mossad, O. et al. Gut microbiota drives age-related oxidative stress and mitochondrial damage in microglia via the metabolite N6-carboxymethyllysine. Nat Neurosci 25, 295–305 (2022).

17. Erny, D. et al. Microbiota-derived acetate enables the metabolic fitness of the brain innate immune system during health and disease. Cell Metab 33, 2260–2276.e7 (2021).

18. Blacher, E. et al. Potential roles of gut microbiome and metabolites in modulating ALS in mice. Nature 572, 474–480 (2019).

19. Sampson, T. R. et al. Gut Microbiota Regulate Motor Deficits and Neuroinflammation in a Model of Parkinson’s Disease. Cell 167, 1469–1480.e12 (2016).

20. van Kessel, S. P. & El Aidy, S. Bacterial Metabolites Mirror Altered Gut Microbiota Composition in Patients with Parkinson’s Disease. Journal of Parkinson’s Disease 9, S359–S370 (2019).

21. Romano, S. et al. Meta-analysis of the Parkinson’s disease gut microbiome suggests alterations linked to intestinal inflammation. npj Parkinsons Dis. 7, 27 (2021).

22. Keshavarzian, A. et al. Colonic bacterial composition in Parkinson’s disease. Movement Disorders 30, 1351–1360 (2015).

23. Boktor, J. Integrated multi-cohort analysis of the Parkinson’s disease gut metagenome. Zenodo 10.5281/zenodo.7183678 (2022).

24. Wallen, Z. D. et al. Metagenomics of Parkinson’s disease implicates the gut microbiome in multiple disease mechanisms. Nat Commun 13, 6958 (2022).

25. Bedarf, J. R. et al. Functional implications of microbial and viral gut metagenome changes in early stage L-DOPA-naïve Parkinson’s disease patients. Genome Medicine 9, 39 (2017).

26. Rosario, D. et al. Systematic analysis of gut microbiome reveals the role of bacterial folate and homocysteine metabolism in Parkinson’s disease. Cell Reports 34, 108807 (2021).

27. Maini Rekdal, V., Bess, E. N., Bisanz, J. E., Turnbaugh, P. J. & Balskus, E. P. Discovery and inhibition of an interspecies gut bacterial pathway for Levodopa metabolism. Science 364, eaau6323 (2019).

28. Goldin, B. R., Peppercorn, M. A. & Goldman, P. CONTRIBUTIONS OF HOST AND INTESTINAL MICROFLORA IN THE METABOLISM OF l-DOPA BY THE RAT. J Pharmacol Exp Ther 186, 160–166 (1973).

29. LeWitt, P. A., Li, J., Lu, M., Guo, L. & Auinger, P. Metabolomic biomarkers as strong correlates of Parkinson disease progression. Neurology 88, 862–869 (2017).

30. Sinclair, E. et al. Metabolomics of sebum reveals lipid dysregulation in Parkinson’s disease. Nat Commun 12, 1592 (2021).

31. Hatano, T., Saiki, S., Okuzumi, A., Mohney, R. P. & Hattori, N. Identification of novel biomarkers for Parkinson’s disease by metabolomic technologies. J Neurol Neurosurg Psychiatry 87, 295–301 (2016).

32. Kaddurah-Daouk, R. & Krishnan, K. R. R. Metabolomics: a global biochemical approach to the study of central nervous system diseases. Neuropsychopharmacology 34, 173–186 (2009).

33. Bhinderwala, F. et al. Metabolomics analyses from tissues in parkinson’s disease. in Methods in Molecular Biology 217–257 (Humana Press Inc., 2019). doi:10.1007/978-1-4939-9488-5_19.

34. Wen, M. et al. Serum uric acid levels in patients with Parkinson’s disease: A meta-analysis. PLOS ONE 12, e0173731 (2017).

35. Bar, N. et al. A reference map of potential determinants for the human serum metabolome. Nature 588, 135–140 (2020).

36. Rockenstein, E. et al. Differential neuropathological alterations in transgenic mice expressing α-synuclein from the platelet-derived growth factor and Thy-1 promoters. J of Neuroscience Research 68, 568–578 (2002).

37. Chesselet, M.-F. et al. A Progressive Mouse Model of Parkinson’s Disease: The Thy1-aSyn (“Line 61”) Mice. Neurotherapeutics 9, 297–314 (2012).

38. Liang, D., et al. *Escherichia coli* triggers α-synuclein pathology in the *LRRK2* transgenic mouse model of PD. Gut Microbes 15, 2276296 (2023).

39. Matheoud, D. et al. Intestinal infection triggers Parkinson’s disease-like symptoms in Pink1−/− mice. Nature 571, 565–569 (2019).

40. Gil-Martinez, A. L. et al. Identification of differentially expressed genes profiles in a combined mouse model of Parkinsonism and colitis. Sci Rep 10, 13147 (2020).

41. Houser, M. C. et al. Experimental colitis promotes sustained, sex-dependent, T-cell-associated neuroinflammation and parkinsonian neuropathology. acta neuropathol commun 9, 139 (2021).

42. Abdel-Haq, R. et al. A prebiotic diet modulates microglial states and motor deficits in α-synuclein overexpressing mice. eLife 11, e81453 (2022).

43. Yan, Y. et al. Gut microbiota and metabolites of α-synuclein transgenic monkey models with early stage of Parkinson’s disease. NPJ Biofilms Microbiomes 7, 69 (2021).

44. Joers, V. et al. Microglia, inflammation and gut microbiota responses in a progressive monkey model of Parkinson’s disease: A case series. Neurobiology of Disease 144, 105027 (2020).

45. Boktor, J. C. et al. Integrated Multi-Cohort Analysis of the Parkinson’s Disease Gut Metagenome. Movement Disorders 38, 399–409 (2023).

46. Cirstea, M. S. et al. Microbiota Composition and Metabolism Are Associated With Gut Function in Parkinson’s Disease. Movement Disorders 35, 1208–1217 (2020).

47. Hertel, J. et al. Integrated Analyses of Microbiome and Longitudinal Metabolome Data Reveal Microbial-Host Interactions on Sulfur Metabolism in Parkinson’s Disease. Cell Reports 29, 1767–1777.e8 (2019).

48. Hasegawa, S. et al. Intestinal Dysbiosis and Lowered Serum Lipopolysaccharide-Binding Protein in Parkinson’s Disease. PLoS ONE 10, e0142164 (2015).

49. Hertel, J. et al. Integrated Analyses of Microbiome and Longitudinal Metabolome Data Reveal Microbial-Host Interactions on Sulfur Metabolism in Parkinson’s Disease. Cell Reports 29, 1767–1777.e8 (2019).

50. Cuvelier, E. et al. Overexpression of Wild-Type Human Alpha-Synuclein Causes Metabolism Abnormalities in Thy1-aSYN Transgenic Mice. Front Mol Neurosci 11, 321 (2018).

51. Videlock, E. J. et al. Distinct Patterns of Gene Expression Changes in the Colon and Striatum of Young Mice Overexpressing Alpha-Synuclein Support Parkinson’s Disease as a Multi-System Process. JPD 13, 1127–1147 (2023).

52. Wrasidlo, W. et al. A *de novo* compound targeting α-synuclein improves deficits in models of Parkinson’s disease. Brain 139, 3217–3236 (2016).

53. Subramaniam, S. R. et al. Chronic nicotine improves cognitive and social impairment in mice overexpressing wild type α-synuclein. Neurobiology of Disease 117, 170–180 (2018).

54. Van Der Veen, J. N. et al. The critical role of phosphatidylcholine and phosphatidylethanolamine metabolism in health and disease. Biochimica et Biophysica Acta (BBA) - Biomembranes 1859, 1558–1572 (2017).

55. Xing, C. et al. Roles of bile acids signaling in neuromodulation under physiological and pathological conditions. Cell Biosci 13, 106 (2023).

56. Heianza, Y. et al. Changes in bile acid subtypes and improvements in lipid metabolism and atherosclerotic cardiovascular disease risk: the Preventing Overweight Using Novel Dietary Strategies (POUNDS Lost) trial. The American Journal of Clinical Nutrition S0002916524001655 (2024) doi:10.1016/j.ajcnut.2024.02.019.

57. Wilson, A., Almousa, A., Teft, W. A. & Kim, R. B. Attenuation of bile acid-mediated FXR and PXR activation in patients with Crohn’s disease. Sci Rep 10, 1866 (2020).

58. MahmoudianDehkordi, S. et al. Altered bile acid profile associates with cognitive impairment in Alzheimer’s disease—An emerging role for gut microbiome. Alzheimer’s & Dementia 15, 76–92 (2019).

59. Baloni, P. et al. Metabolic Network Analysis Reveals Altered Bile Acid Synthesis and Metabolism in Alzheimer’s Disease. Cell Reports Medicine 1, 100138 (2020).

60. Varma, V. R. et al. Bile acid synthesis, modulation, and dementia: A metabolomic, transcriptomic, and pharmacoepidemiologic study. PLoS Med 18, e1003615 (2021).

61. Karbowska, M. et al. Neurobehavioral effects of uremic toxin-indoxyl sulfate in the rat model. Sci Rep 10, 9483 (2020).

62. Sun, C.-Y. et al. Indoxyl sulfate caused behavioral abnormality and neurodegeneration in mice with unilateral nephrectomy. Aging (Albany NY) 13, 6681–6701 (2021).

63. Kalia, L. V. & Lang, A. E. Parkinson’s disease. Lancet 386, 896–912 (2015).

64. Yao, Y. & Han, W. Proline Metabolism in Neurological and Psychiatric Disorders. Mol.Cells 45, 781–788 (2022).

65. Bruce, K. D., Zsombok, A. & Eckel, R. H. Lipid Processing in the Brain: A Key Regulator of Systemic Metabolism. Front. Endocrinol. 8, (2017).

66. Yoneshiro, T. et al. BCAA catabolism in brown fat controls energy homeostasis through SLC25A44. Nature 572, 614–619 (2019).

67. Hai, X. et al. Mechanism of Prominent Trimethylamine Oxide (TMAO) Accumulation in Hemodialysis Patients. PLoS ONE 10, e0143731 (2015).

68. Wang, Z. et al. Non-lethal Inhibition of Gut Microbial Trimethylamine Production for the Treatment of Atherosclerosis. Cell 163, 1585–1595 (2015).

69. Nie, J. et al. Serum Trimethylamine N-Oxide Concentration Is Positively Associated With First Stroke in Hypertensive Patients. Stroke 49, 2021–2028 (2018).

70. Manor, O. et al. A Multi-omic Association Study of Trimethylamine N-Oxide. Cell Reports 24, 935–946 (2018).

71. Dumas, M.-E. et al. Microbial-Host Co-metabolites Are Prodromal Markers Predicting Phenotypic Heterogeneity in Behavior, Obesity, and Impaired Glucose Tolerance. Cell Reports 20, 136–148 (2017).

72. Dehghan, P., Farhangi, M. A., Nikniaz, L., Nikniaz, Z. & Asghari-Jafarabadi, M. Gut microbiota-derived metabolite trimethylamine N-oxide (TMAO) potentially increases the risk of obesity in adults: An exploratory systematic review and dose-response meta-analysis. Obesity Reviews 21, e12993 (2020).

73. Sankowski, B. et al. Higher cerebrospinal fluid to plasma ratio of p-cresol sulfate and indoxyl sulfate in patients with Parkinson’s disease. Clinica Chimica Acta 501, 165–173 (2020).

74. Chen, S. et al. The Gut Metabolite Trimethylamine N-oxide Is Associated With Parkinson’s Disease Severity and Progression. Movement Disorders 35, 2115–2116 (2020).

75. Zhou, H., et al. Causal effect of gut-microbiota-derived metabolite trimethylamine N-oxide on Parkinson’s disease: A Mendelian randomization study. Euro J of Neurology 30, 3451–3461 (2023).

76. Chung, S. J. et al. Gut microbiota-derived metabolite trimethylamine N-oxide as a biomarker in early Parkinson’s disease. Nutrition 83, 111090 (2021).

77. Fanning, S., Selkoe, D. & Dettmer, U. Parkinson’s disease: proteinopathy or lipidopathy? npj Parkinsons Dis. 6, 3 (2020).

78. Whiley, L. et al. Evidence of altered phosphatidylcholine metabolism in Alzheimer’s disease. Neurobiology of Aging 35, 271–278 (2014).

79. Cooper, O., Hallett, P. & Isacson, O. Upstream lipid and metabolic systems are potential causes of Alzheimer’s disease, Parkinson’s disease and dementias. The FEBS Journal febs.16638 (2022) doi:10.1111/febs.16638.

80. O’Leary, E. I., Jiang, Z., Strub, M.-P. & Lee, J. C. Effects of phosphatidylcholine membrane fluidity on the conformation and aggregation of N-terminally acetylated α-synuclein. Journal of Biological Chemistry 293, 11195–11205 (2018).

81. Cheng, D. et al. Lipid Pathway Alterations in Parkinson’s Disease Primary Visual Cortex. PLoS ONE 6, e17299 (2011).

82. Farmer, K., Smith, C., Hayley, S. & Smith, J. Major Alterations of Phosphatidylcholine and Lysophosphotidylcholine Lipids in the Substantia Nigra Using an Early Stage Model of Parkinson’s Disease. IJMS 16, 18865–18877 (2015).

83. Korytowski, A. et al. Accumulation of phosphatidylcholine on gut mucosal surface is not dominated by electrostatic interactions. Biochimica et Biophysica Acta (BBA) - Biomembranes 1859, 959–965 (2017).

84. Warnecke, T., Schäfer, K.-H., Claus, I., Del Tredici, K. & Jost, W. H. Gastrointestinal involvement in Parkinson’s disease: pathophysiology, diagnosis, and management. npj Parkinsons Dis. 8, 31 (2022).

85. Miletić Vukajlović, J., et al. Increased plasma phosphatidylcholine/lysophosphatidylcholine ratios in patients with Parkinson’s disease. Rapid Comm Mass Spectrometry 34, e8595 (2020).

86. Guerreiro, P. S. et al. Mutant A53T α-Synuclein Improves Rotarod Performance Before Motor Deficits and Affects Metabolic Pathways. Neuromolecular Med 19, 113–121 (2017).

87. Zhang, L. et al. Circulating Cholesterol Levels May Link to the Factors Influencing Parkinson’s Risk. Front Neurol 8, 501 (2017).

88. Guo, X. et al. The serum lipid profile of Parkinson’s disease patients: a study from China. Int J Neurosci 125, 838–844 (2015).

89. Fu, X. et al. A systematic review and meta-analysis of serum cholesterol and triglyceride levels in patients with Parkinson’s disease. Lipids in Health and Disease 19, 97 (2020).

90. Barupal, D. K. et al. Sets of coregulated serum lipids are associated with Alzheimer’s disease pathophysiology. Alz & Dem Diag Ass & Dis Mo 11, 619–627 (2019).

91. Hamilton, L. K. et al. Aberrant Lipid Metabolism in the Forebrain Niche Suppresses Adult Neural Stem Cell Proliferation in an Animal Model of Alzheimer’s Disease. Cell Stem Cell 17, 397–411 (2015).

92. Yin, F. Lipid metabolism and Alzheimer’s disease: clinical evidence, mechanistic link and therapeutic promise. The FEBS Journal 290, 1420–1453 (2023).

93. Li, P. et al. Gut Microbiota Dysbiosis Is Associated with Elevated Bile Acids in Parkinson’s Disease. Metabolites 11, 29 (2021).

94. Graham, S. F. et al. Metabolomic Profiling of Bile Acids in an Experimental Model of Prodromal Parkinson’s Disease. Metabolites 8, E71 (2018).

95. Rosario, D., et al. Systematic analysis of gut microbiome reveals the role of bacterial folate and homocysteine metabolism in Parkinson’s disease. Cell Reports 34, (2021).

96. Lin, Y.-T. et al. Indoxyl Sulfate Induces Apoptosis Through Oxidative Stress and Mitogen-Activated Protein Kinase Signaling Pathway Inhibition in Human Astrocytes. JCM 8, 191 (2019).

97. Adesso, S. et al. Indoxyl Sulfate Affects Glial Function Increasing Oxidative Stress and Neuroinflammation in Chronic Kidney Disease: Interaction between Astrocytes and Microglia. Front. Pharmacol. 8, 370 (2017).

98. Brydges, C. R. et al. Indoxyl sulfate, a gut microbiome-derived uremic toxin, is associated with psychic anxiety and its functional magnetic resonance imaging-based neurologic signature. Sci Rep 11, 21011 (2021).

99. Watanabe, K. et al. Effect of uremic toxins on hippocampal cell damage: analysis in vitro and in rat model of chronic kidney disease. Heliyon 7, e06221 (2021).

100. Cassani, E. et al. Increased urinary indoxyl sulfate (indican): New insights into gut dysbiosis in Parkinson’s disease. Parkinsonism & Related Disorders 21, 389–393 (2015).

101. Wu, G. Important roles of dietary taurine, creatine, carnosine, anserine and 4-hydroxyproline in human nutrition and health. Amino Acids 52, 329–360 (2020).

102. Chen, X., Xie, C., Sun, L., Ding, J. & Cai, H. Longitudinal Metabolomics Profiling of Parkinson’s Disease-Related α-Synuclein A53T Transgenic Mice. PLoS ONE 10, e0136612 (2015).

103. Picca, A. et al. Circulating amino acid signature in older people with Parkinson’s disease: A metabolic complement to the EXosomes in PArkiNson Disease (EXPAND) study. Experimental Gerontology 128, 110766 (2019).

104. Shao, Y. et al. Comprehensive metabolic profiling of Parkinson’s disease by liquid chromatography-mass spectrometry. Mol Neurodegeneration 16, 4 (2021).

105. Gonzalez-Riano, C. et al. Prognostic biomarkers of Parkinson’s disease in the Spanish EPIC cohort: a multiplatform metabolomics approach. npj Parkinsons Dis. 7, 73 (2021).

106. O’Mahony, S. M., Clarke, G., Borre, Y. E., Dinan, T. G. & Cryan, J. F. Serotonin, tryptophan metabolism and the brain-gut-microbiome axis. Behavioural Brain Research 277, 32–48 (2015).

107. Lou, H. Dopamine precursors and brain function in phenylalanine hydroxylase deficiency. Acta Paediatrica 83, 86–88 (1994).

108. Havelund, J., Heegaard, N., Færgeman, N. & Gramsbergen, J. Biomarker Research in Parkinson’s Disease Using Metabolite Profiling. Metabolites 7, 42 (2017).

109. Peng, W., Schröder, L. F., Song, P., Wong, Y. C. & Krainc, D. Parkin regulates amino acid homeostasis at mitochondria-lysosome (M/L) contact sites in Parkinson’s disease. Sci. Adv. 9, eadh3347 (2023).

110. Yue, C. et al. Trimethylamine N-oxide prime NLRP3 inflammasome via inhibiting ATG16L1-induced autophagy in colonic epithelial cells. Biochemical and Biophysical Research Communications 490, 541–551 (2017).

111. Lee, Y. et al. Longitudinal Plasma Measures of Trimethylamine N-Oxide and Risk of Atherosclerotic Cardiovascular Disease Events in Community-Based Older Adults. Journal of the American Heart Association 10, e020646 (2021).

112. Vogt, N. M. et al. The gut microbiota-derived metabolite trimethylamine N-oxide is elevated in Alzheimer’s disease. Alzheimer’s Research & Therapy 10, 124 (2018).

113. Li, D. et al. Trimethylamine-*N* -oxide promotes brain aging and cognitive impairment in mice. Aging Cell 17, e12768 (2018).

114. Hoyles, L. et al. Regulation of blood–brain barrier integrity by microbiome-associated methylamines and cognition by trimethylamine N-oxide. Microbiome 9, 235 (2021).

115. Brunt, V. E. et al. The gut microbiome–derived metabolite trimethylamine N-oxide modulates neuroinflammation and cognitive function with aging. GeroScience 43, 377–394 (2021).

116. Kalecký, K. & Bottiglieri, T. Targeted metabolomic analysis in Parkinson’s disease brain frontal cortex and putamen with relation to cognitive impairment. npj Parkinsons Dis. 9, 84 (2023).

117. Chen, S.-J. et al. The Gut Metabolite Trimethylamine N-oxide Is Associated With Parkinson’s Disease Severity and Progression. Mov Disord 35, 2115–2116 (2020).

118. Ferreon, A. C. M., Moosa, M. M., Gambin, Y. & Deniz, A. A. Counteracting chemical chaperone effects on the single-molecule α-synuclein structural landscape. Proceedings of the National Academy of Sciences 109, 17826–17831 (2012).

119. Jamal, S., Kumari, A., Singh, A., Goyal, S. & Grover, A. Conformational Ensembles of α-Synuclein Derived Peptide with Different Osmolytes from Temperature Replica Exchange Sampling. Frontiers in Neuroscience 11, (2017).

120. Uversky, V. N., Li, J. & Fink, A. L. Trimethylamine-N-oxide-induced folding of alpha-synuclein. FEBS Lett 509, 31–35 (2001).

121. Chesselet, M.-F. et al. A Progressive Mouse Model of Parkinson’s Disease: The Thy1-aSyn (“Line 61”) Mice. Neurotherapeutics 9, 297–314 (2012).

122. Needham, B. D. et al. A gut-derived metabolite alters brain activity and anxiety behaviour in mice. Nature 602, 647–653 (2022).

123. Paxinos, G. & Franklin, K. B. J. Paxinos and Franklin’s The Mouse Brain in Stereotaxic Coordinates. (Elsevier, Academic Press, London San Diego Cambridge; MA Kidlington, Oxford, 2019).

124. St John-Williams, L., et al. Targeted metabolomics and medication classification data from participants in the ADNI1 cohort. Sci Data 4, 170140 (2017).

125. St John-Williams, L., et al. Bile acids targeted metabolomics and medication classification data in the ADNI1 and ADNIGO/2 cohorts. Sci Data 6, 212 (2019).

